# Positive cooperativity between RAS-binding and cysteine-rich domains regulates RAF membrane binding kinetics via lateral rebinding

**DOI:** 10.1101/2025.04.23.650295

**Authors:** Andres Jimenez Salinas, Kesaria Tevdorashvili, Julian Grim, Alexia Morales, Ani Chakhrakia, Young Kwang Lee

## Abstract

RAF activation requires interactions with both RAS nanoclusters and membrane lipids, yet the molecular basis of this process remains unclear. Using a bottom-up reconstitution approach, we show how coordinated protein–protein and protein–lipid interactions regulate membrane binding dynamics of RAF to drive its multistep activation. Within membrane environments, the RAS-binding domain (RBD) and cysteine-rich domain (CRD) exhibit cooperativity, with CRD-mediated phosphatidylserine binding stabilizing the RBD:RAS complex. Importantly, RAF remains membrane-bound through lateral rebinding to RAS, where a weak CRD–lipid interaction plays an essential role. This lateral rebinding extends RAF’s membrane dwell time under high RAS density conditions, which are found in RAS nanoclusters. This prolonged membrane residence likely facilitates kinetic proofreading of RAF’s multistep activation within RAS nanoclusters, ensuring signaling specificity. Given the high abundance of weak multivalent membrane interactions, lateral rebinding may be a common mechanism for regulating the activity of signaling proteins through sustained membrane retention.

## Background

The RAF kinases are critical regulators of the mitogen-activated protein kinase (MAPK) pathway, controlling diverse cellular processes such as proliferation, differentiation, and survival^1–3^. RAF isoforms share a conserved domain organization: an N-terminal regulatory region composed of the RAS-binding domain (RBD) and the cysteine-rich domain (CRD), and a C-terminal kinase domain (Fig. 1a). In their basal state, RAF kinases reside in the cytosol as autoinhibited monomers, and they are activated when the RBD binds to GTP-loaded RAS at the membrane surface. Once activated, RAF initiates the three-tiered RAF-MEK-ERK kinase cascade, serving as a central node that modulates pathway dynamics. Mutations in RAF can lead to constitutive signaling in cancers and developmental disorders^4–7^. Paradoxical activation has been observed when inhibitors targeting oncogenic BRAF^V600E^ are used under high RAS·GTP conditions, inadvertently promoting RAF dimerization and unintended ERK pathway activation^8–14^. Additionally, drugs that enhance RAF dimerization can alter the RAS-ERK signal transmission kinetics, resulting in improper cellular behavior, such as hyperproliferation^15^.

**Fig. 1:**
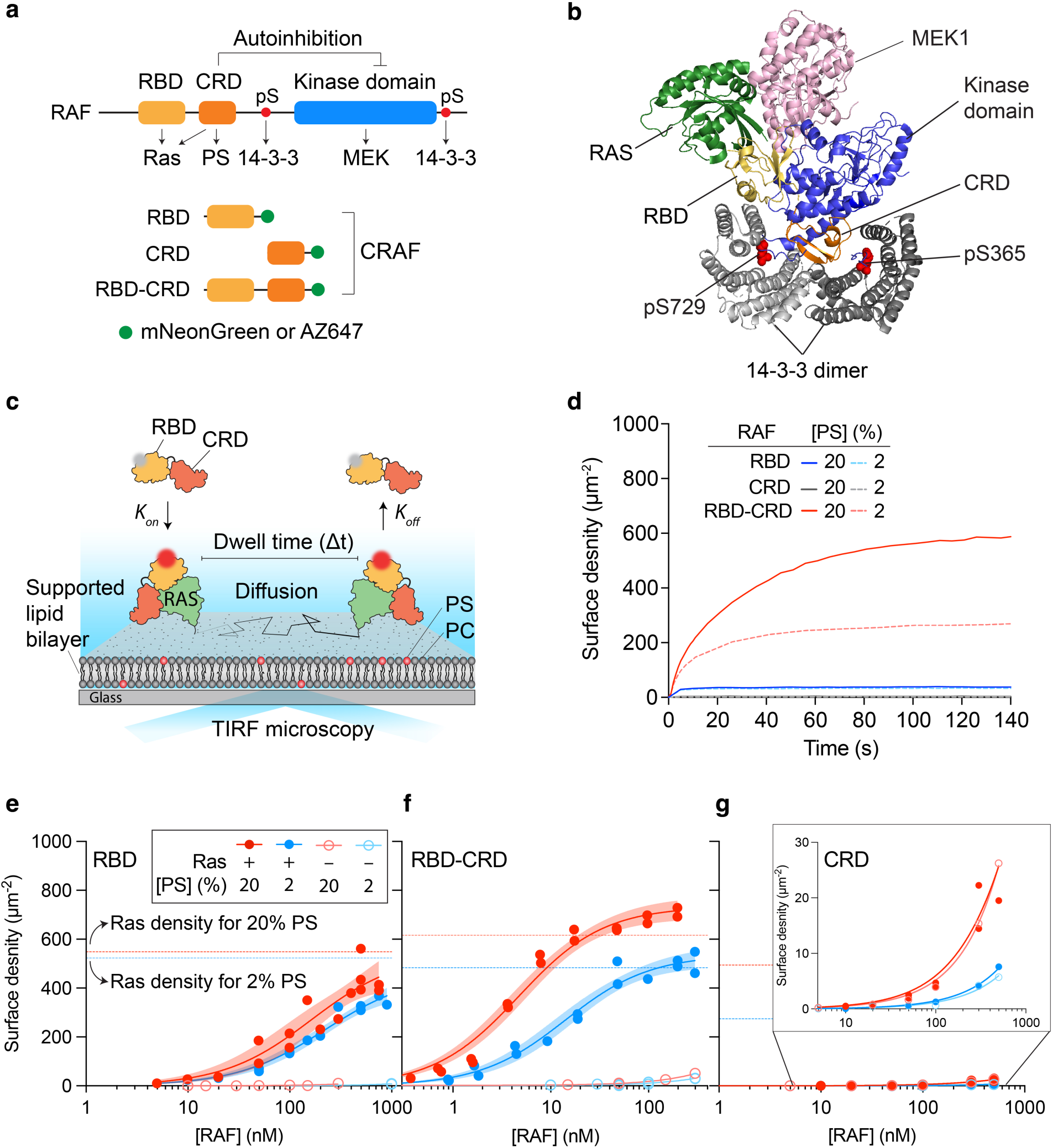
RBD-dependent lipid engagement of CRD induces synergistic binding on RAS-functionalized supported lipid bilayers. **a** Domain diagrams of RAF kinase and CRAF N-terminal regulatory domain constructs used in this study. **b** Structure of the KRAS/BRAF/MEK1/14-3-3 complex (PDB:8DGS). In the absence of a membrane environment, BRAF remains autoinhibited in complex with KRAS. **c** Schematic illustration of RAF binding assays on RAS-functionalized supported lipid bilayers using TIRF microscopy. Fluorescent labels are represented as a grey circle (outside the TIRF volume) and a red circle (within the TIRF volume). **d** Binding kinetics of different CRAF constructs on RAS-functionalized membranes containing 2% or 20% PS. RAF concentration: 20 nM. RAS surface density: ∼600 μm^-2^. **e-g** RAF titration binding curves, with the shaded regions representing the 95% confidence range. Apparent dissociation constant, *K*_d_, ± 95% confidence interval on RAS-functionalized membranes was determined to be 222 ± 88 (2% PS) and 163 ± 126 (20% PS) for RBD (e), and 14 ± 5 (2% PS) and 5 ± 1 (20% PS) for RBD-CRD (f). For all other conditions, *K*_d_ was too high to be determined using TIRF microscopy. RAS surface densities are indicated with dashed lines. *K*_d_ values were estimated by fitting a one-site specific binding model.

RAF activation proceeds through multiple intermediates governed by complex molecular interactions^1^. However, the precise mechanisms underlying the transition from inactive monomer to active dimer remain to be fully determined. Both cellular and *in vitro* studies highlight the critical role of the membrane environment in this process^18–21^. In the active state, RAF associates with the plasma membrane via interaction of its RBD with active RAS×GTP^21–24^. The CRD of RAF further stabilizes its membrane association by engaging both RAS and negatively charged lipids, such as phosphatidylserine (PS)^25–30^. Thus, anchored at the membrane, RAF becomes dephosphorylated by the MRAS-SHOC2-PP1C phosphatase complex and dimerizes via its kinase domain—an essential step for allosteric activation^31–35^. The 14-3-3 dimer reinforces an active dimeric assembly by binding a phosphoserine-containing motif on each RAF kinase domain^36–41^.

Recent cryo-electron microscopy (EM) structures of BRAF provide important insight into how RAF activity is regulated^39,40^. In its inactive monomeric form, BRAF forms an autoinhibited complex with MEK1 and a 14-3-3 dimer (Fig. 1b). The 14-3-3 dimer binds two phosphoserine residues within the CR2 and CR3 regions of the same RAF molecule, forming a cradle-like structure that sequesters the CRD. The CRD is a critical component that stabilizes the overall autoinhibited complex by making multiple contacts with the kinase domain and 14-3-3. The autoinhibited BRAF structure suggests that extraction of the CRD following RBD binding to RAS could destabilize the autoinhibited complex, promoting the transition from a monomeric to a dimeric state. However, the precise mechanism requires further investigation. A cryo-EM RAS:RAF recruitment complex structure revealed that RAS binding to the RBD alone is insufficient to release RAF from autoinhibition, emphasizing the critical role of coordinated protein and lipid interactions^42^.

A key aspect of RAF activation is that Ras nanoclusters serve as the primary sites for recruitment and activation of RAF^43–45^. RAS nanoclusters locally enrich specific lipids that support RAF activation, underscoring the importance of integrating protein and lipid inputs^2^. The RBD and CRD—responsible for binding protein and lipids, respectively—are arranged in tandem and separated by a short, six-amino-acid-long linker. X-ray crystal structures show that the RBD and CRD together form an extended structural unit when bound to RAS^46,47^, yet in solution the CRD contributes little to RAS binding^47^. It is not known whether the RBD and CRD mutually modify their respective binding interactions within membrane environments. The membrane binding mechanisms and kinetics arising from their functional coupling also remain unknown—particularly how these two tandem domains integrate protein and lipid interactions within RAS nanoclusters to drive multistep activation of RAF.

Here, we employed a bottom-up reconstitution approach with supported lipid bilayers (SLBs) to systematically investigate how RAS proteins and lipids cooperatively regulate RAF membrane binding dynamics. By quantifying the kinetics, affinities, and diffusion of various CRAF regulatory domain constructs on RAS-functionalized membranes, we demonstrate that the RBD and CRD functionally influence each other within membrane environments.

Specifically, RBD binding to RAS facilitates engagement of the RAF CRD with negatively charged PS lipids, resulting in two kinetically distinct steps during the binding of protein and lipid partners. This sequential association enables the CRD to bind abundant PS lipids in a RAS·GTP-dependent manner, ensuring that RAF membrane binding is strictly initiated by RAS activation. Moreover, CRD–lipid engagement reinforces RBD–RAS association, effectively reducing the membrane dissociation rate. Importantly, we found that RAF maintains its membrane-bound state through lateral rebinding to RAS, with a weak CRD–lipid interaction playing an essential role. This lateral rebinding mechanism significantly extends RAF’s membrane dwell time under conditions of high RAS density, a defining characteristic of RAS nanoclusters. RAS nanoclusters may facilitate kinetic proofreading of RAF activation by increasing the likelihood of completing the multistep activation sequence through prolonging membrane residence. Overall, our study offers new insight into how kinetic regulation by coordinated protein and lipid inputs within RAS nanoclusters modulates RAF signaling, advancing our understanding of signaling activation and specificity.

## Results

### RBD binding to RAS promotes lipid engagement of CRD

To investigate the mechanisms underlying RAF membrane binding, we reconstituted CRAF and RAS interaction on a SLB (Fig. 1a,c). HRAS (1-184, C118S, hereafter referred to as RAS), which retains its native cysteine palmitoylation sites (181 and 184) in the hypervariable region, was linked to lipids via maleimide chemistry. This widely used lipidation method preserves the biological function and structure of RAS and has been applied to all three RAS isoforms^48–52^.

RAS was then activated by exchanging nucleotides with GTP using the catalytic domain of Son of Sevenless (SOS) (Supplementary Fig. 1)^49^. The membrane binding kinetics of fluorescently labeled CRAF constructs (mNeonGreen or AZ647) were measured using total internal reflection fluorescence (TIRF) microscopy (Fig. 1c). The surface density of membrane-bound CRAF was determined from TIRF intensity, calibrated by fluorescence correlation spectroscopy (FCS) (Supplementary Fig. 2)^53^.

RAF is a low abundance protein, existing in the cytoplasm at a concentration ranging from high picomolar to low nanomolar^54^. Accordingly, we characterized membrane binding kinetics of truncated CRAF regulatory domains within this physiologically relevant range. At 20 nM, the RBD exhibited low-level, fast equilibrium binding (∼30 molecules per µm^2^). The binding was unaffected by PS content (2% vs 20) (blue, Fig. 1d). Despite the known affinity of the CRD for PS, 20 nM of CRD displayed insignificant binding (∼1 to 2 molecules per µm^2^) on both 2% and 20% PS membranes functionalized with RAS (gray, Fig. 1d). Notably, the tandem RBD-CRD exhibited a marked increase in membrane binding (∼600 molecules per µm^2^) that depended on PS and had slower kinetics (red, Fig. 1d). This finding is intriguing because the CRD alone shows negligible binding, yet it effectively engages PS lipids when arranged in tandem with RBD. To test whether the enhanced lipid binding of the CRD requires active RAS and RBD interactions, we measured the binding of RBD-CRD on PS membranes functionalized with inactive RAS·GDP. Indeed, negligible binding was observed, indicating that the enhanced lipid sensing exhibited by CRD requires a specific interaction between RAS·GTP and the RBD (Supplementary Fig. 3). Collectively, these results demonstrate that the RBD and CRD are functionally coupled, such that RBD binding to RAS promotes the CRD for lipid engagement.

To better understand how RAF integrates protein and lipid inputs for membrane binding, we measured CRAF affinity to RAS-free and RAS-functionalized membranes with different PS concentrations. The isolated RBD exhibited apparent dissociation constants (*K_d_*) that depended little upon PS content, measuring 222 nM and 163 nM on RAS-tethered membranes with 2% and 20% PS, respectively (Fig. 1e). These values are consistent with previously reported affinities of the CRAF RBD for HRAS determined independent of membranes^47,55^. Without RAS, RBD binding was minimal at all concentrations tested. In contrast, RBD-CRD exhibited an affinity of more than an order of magnitude higher than that of the RBD alone (Fig. 1f). Specifically, the *K_d_* for RBD-CRD was determined to be 14 nM and 5 nM on RAS-functionalized membranes containing 2% and 20% PS, respectively. The evident dependence on PS concentration for RBD-CRD confirms that the augmented membrane binding is driven by CRD and PS lipid interaction. Notably, the affinity of RBD-CRD for RAS on membranes, as determined here, is significantly higher than that previously measured in non-membrane environments (∼150 nM, similar to isolated RBD)^47^. This difference underscores an indispensable role of weak CRD–lipid interaction in achieving high-affinity membrane association. Removing RAS essentially abolished this high-affinity binding, affirming that lipid engagement by the CRD depends on RBD and RAS interaction. (Fig. 1f). The CRD binds RAS with micromolar-scale binding affinity via an interface distinct from that employed by RAS in binding to the RBD^26^. The observed weak interaction between RAS and isolated CRD, lacking RBD, does not independently facilitate lipid engagement by the CRD. As shown in Figure 1g, the isolated CRD displayed similarly very low levels of binding on both RAS-free and RAS-functionalized membranes.

By quantifying the surface density of the proteins, we found that the density of RBD-CRD at the membrane was higher than that of RAS (Fig. 1f). This indicates the presence of two populations of RBD-CRD: one associated with RAS and the other not. Notably, the recruitment of excess RBD-CRD, which is associated only with lipids, is still largely mediated by the RAS·GTP interaction, as the excess RBD-CRD density was higher than that observed on RAS-free membranes. The existence of RBD-CRD in two distinct membrane-bound states forms a fundamental basis for the kinetic control of RAF–membrane interactions through a lateral rebinding mechanism, as will be discussed later in this study.

### CRD and lipid interaction selectively modulates the membrane dissociation rate

Differences in membrane-binding affinities among CRAF regulatory domains must reflect changes in kinetic rate constants, namely membrane association (*k_on_*) and dissociation (*k_off_*) rates. Characterizing these kinetic parameters is important for understanding the mechanisms that govern the initiation and completion of RAF activation. Given that the RBD and CRD have distinct primary binding partners—RAS and lipids, respectively—we first examined how these interactions interplay to modulate *k_on_* and *k_off_.* The *k_on_* values were quantified by counting the single-molecule recruitment events of AZ647-labeled CRAF to the membrane using TIRF microscopy (Supplementary Fig. 4). In line with our ensemble binding measurements, efficient membrane recruitment required the interaction between RBD and RAS (Fig. 2a). Notably, *k_on_* values for both RBD and RBD-CRD were unaffected by PS concentration (Fig. 2b). This suggests that the initial recruitment is primarily driven by the protein–protein interaction between RBD and RAS, without involving CRD and lipid interactions. The CRD exerts a modest inhibitory effect on RBD–RAS binding, reducing the *k_on_* values for RBD-CRD. Because the RBD-CRD exhibits a lower *k_on_* than the RBD, its higher binding affinity must stem from a reduced membrane dissociation rate. We quantified the apparent *k_off_* using an ensemble free-dissociation assay under continuous buffer flow. Indeed, on RAS-functionalized membranes containing 20% PS lipids, mNG-fused RBD-CRD displayed an extended membrane residence time, with a half-life of ∼77 seconds—almost two orders of magnitude longer than the ∼1 second half-life observed for RBD (Fig. 2c). This result indicates that the CRD and PS interaction selectively regulates membrane dissociation kinetics without affecting the association rate.

**Fig 2:**
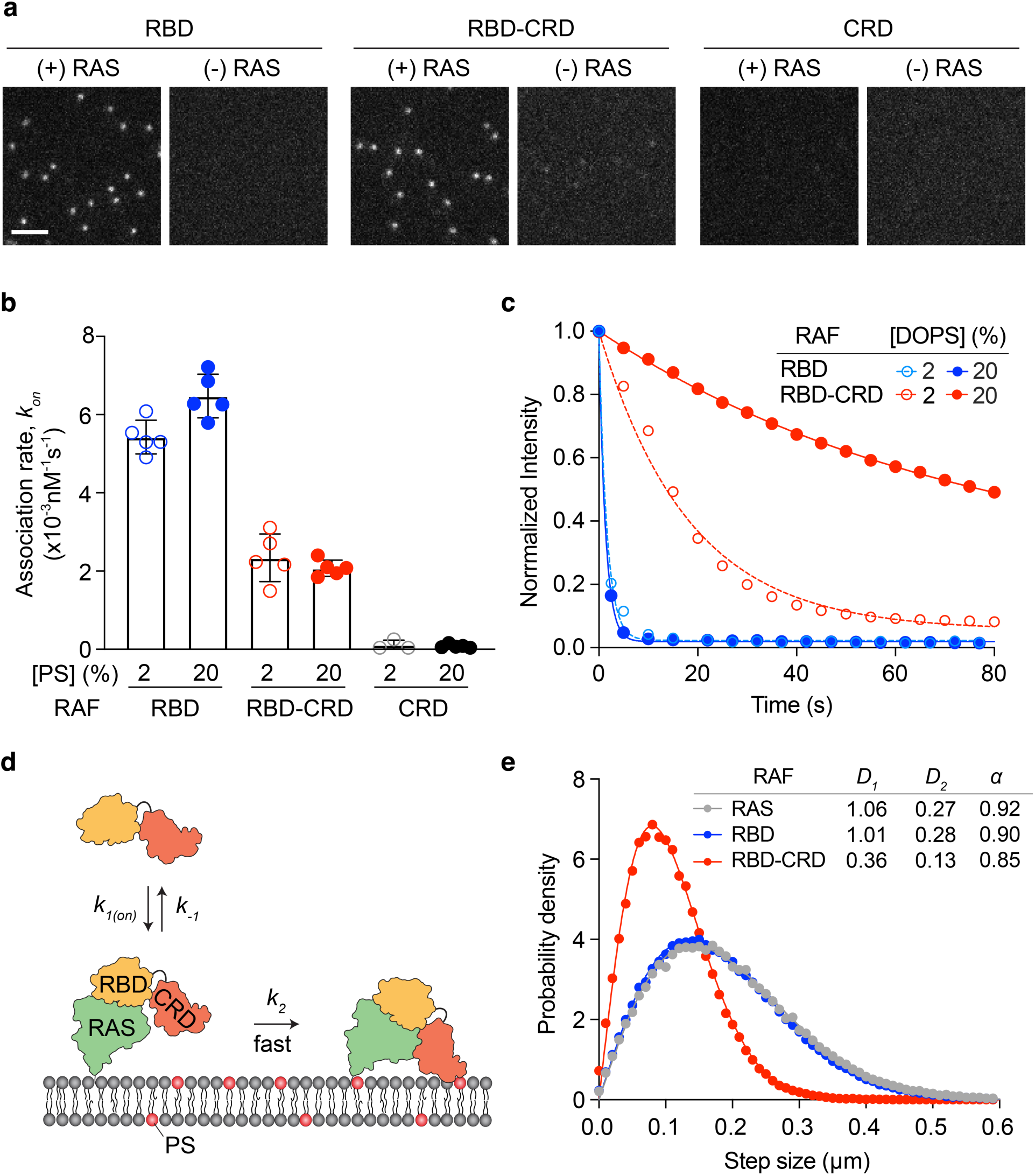
Kinetic analysis reveals that CRD selectively regulates RAF’s membrane dissociation rate. **a** Representative images showing single-molecule association events of RAF constructs on RAS-functionalized and RAS-free membranes. **b** Association rates of various RAF constructs on RAS-functionalized membranes containing 2% or 20% PS. The error bars represent the standard deviations. **c** Ensemble free dissociation curves for RAF constructs on RAS-functionalized membranes containing 2% or 20% PS. The curves were fit to a single-exponential decay to estimate half-lives, while the underlying dissociation mechanism is more complex as discussed later in this study. The half-life is approximately 1 second for RBD at both 2% and 20% PS, and 14 and 77 seconds for RBD-CRD at 2% and 20% PS, respectively. **d** Schematic illustration of two-step membrane association mechanism in which RBD–RAS binding promotes CRD–lipid engagement. **e** Step size distribution analysis of RAS, RBD and RBD-CRD on RAS-functionalized membranes with 20% PS. *D*_1_ and *D*_2_ represent diffusion coefficients of fast and slow species, and α denotes the population of the fast species. Apparent diffusion coefficients (μm^2^/s), calculated by the weighted average of *D*_1_ and *D*_2_, are 1.00 for RAS, 0.94 for RBD, and 0.33 for RBD–CRD. The time interval between frames was 10 ms. RAS surface density (μm^-2^): ∼400 for (a) and (b); ∼500 for (c); ∼550 for (e).

Combined with the affinity measurements, these kinetic analyses suggest that CRAF localizes to membranes through two kinetically distinct steps (Fig. 2d). In the first step, the RBD binds active RAS. In the second step, lipid engagement by the CRD—particularly a productive interaction—occurs only after the RBD has bound to RAS. This two-step mechanism ensures Ras·GTP-specific membrane localization of RAF while leveraging abundant PS lipids to prolong membrane dwell time and achieve high-affinity binding.

### CRD persistently engages PS lipids upon RBD and RAS binding

To investigate how the RAS:RAF complex interacts with lipids, we performed diffusion analysis using single-particle tracking. A membrane containing 20% DOPS was functionalized with RAS at a density of ∼550 per µm^2^. For single particle tracking of RAS, a small fraction of RAS was labeled with Alexa647-GppNp (a fluorescent, nonhydrolyzable GTP analog), ensuring well-dispersed single molecules (Supplementary Fig. 5). RAS alone exhibited two distinct diffusion states (gray, Fig. 2e). The majority (∼90% occupancy, α) had a fast diffusion coefficient of 1.06 µm²/s (*D*_1_). The secondary, slower state exhibited a diffusion coefficient of 0.27 µm²/s (*D*_2_). This minor slow state may result from microscale heterogeneity of PS membranes or strong coupling with the solid support, a behavior also observed in fluorescent lipids (Supplementary Fig. 6). We attribute the majority fast state to RAS with minimal lipid interaction. Similarly, AZ647-labeled RBD bound to unlabeled RAS-functionalized membranes showed minimal lipid interaction and mirrored RAS diffusion (blue, Fig. 2). By contrast, the RBD-CRD diffused significantly more slowly due to lipid engagement by CRD (red, Fig. 2e). Two diffusional states were identified at 0.36 µm²/s (85% occupancy) and 0.13 µm²/s. Even the faster state is substantially slower than that of RBD with minimal lipid interaction, suggesting that once recruited by RAS, the RBD-CRD persistently engages PS lipids. Because RAS-RAF complexes were tracked at the single-molecule level and separated by several micrometers, the observed decrease in diffusion coefficient reflects lipid interactions with individual RAS:RBD-CRD complexes, rather than collective behaviors such as clustering. In support of this, the RBD-CRD at saturation density displayed diffusion behavior identical to that at single-molecule density, confirming the absence of clustering in our system (Supplementary Fig. 7). Previous studies have shown that RAS:RAF clustering requires more complex lipid compositions^56,57^. The profound reduction in diffusion coefficient in our study could be explained by multivalent lipid interactions, facilitated by the membrane insertion of interdigitated hydrophobic and positively charged residues within the CRD loop 1 (residues 143-149) and 2 (residues 157-164)^6,57^.

Engagement of the CRD with lipids appears to be a critical step in triggering the structural rearrangement of the autoinhibited RAF complex. To evaluate the kinetics and efficiency of CRD–lipid interaction following RBD binding, we compared the step-size distributions of RBD-CRD at the initial moment of contact (first-step sizes) versus the entire observation period (all-step sizes). The majority of RBD-CRD rapidly engage lipids within the 20 ms time resolution of the experiment, as evidenced by the similarity between these two distributions (Supplementary Fig. 8). This rapid lipid engagement (Forward Step 2 in Fig. 2d) effectively renders the two-step association mechanism unidirectional, with the first step serving as the rate-determining step. However, in the full-length protein, CRD–lipid binding may be significantly slower, because the lipid-interacting residues of the CRD are occluded by 14-3-3 dimers. The rate of this step likely plays an important role in RAF activation and can be modulated by the degree of autoinhibition, which varies among isoforms and is alleviated by pathogenic mutations and inhibitors^6,58^.

### CRD–lipid engagement enhances the affinity of protein–protein interaction in RAS:RAF complex at the membrane

A previous computational study predicted that CRD–lipid interactions reduce fluctuations between RAS and the RBD due to the short linker, thereby potentially increasing their overall association^59^. To test this prediction experimentally, we developed a FRET-based competition assay to monitor RAS–CRAF dissociation kinetics on membranes (Fig. 3a). In this assay, RAS is labeled with the Atto488-GppNp donor, and CRAF (either RBD or RBD–CRD) is tagged with the AZ647 acceptor. The assay begins by forming the CRAF:RAS complex on the membrane, which generates a FRET signal. A high concentration of unlabeled RBD-CRD (3 µM) is introduced to compete with the labeled CRAF (see Supplementary Fig. 9 for the concentration optimization). Upon dissociation of AZ647-CRAF from RAS, unlabeled CRAF occupies RAS, preventing the rebinding of labeled CRAF. The time-dependent change in FRET signal thus specifically tracks CRAF dissociation from RAS, while excluding dissociation of membrane-bound CRAF that is not associated with RAS. Using this assay, we observed that RBD-CRD dissociates from RAS significantly more slowly than RBD alone (Fig. 3b). The presence of PS lipids further accentuates this slowed dissociation for RBD-CRD. These findings indicate that CRD–lipid interactions enhance the binding strength between RAS and the RBD, thereby stabilizing the complex and slowing its dissociation. Consequently, the protein–protein and protein–lipid interactions within the RAF regulatory domains mutually reinforce each other: RBD binding to RAS facilitates CRD–lipid engagement, which in turn strengthens RBD–RAS association (Fig. 3c).

**Fig. 3:**
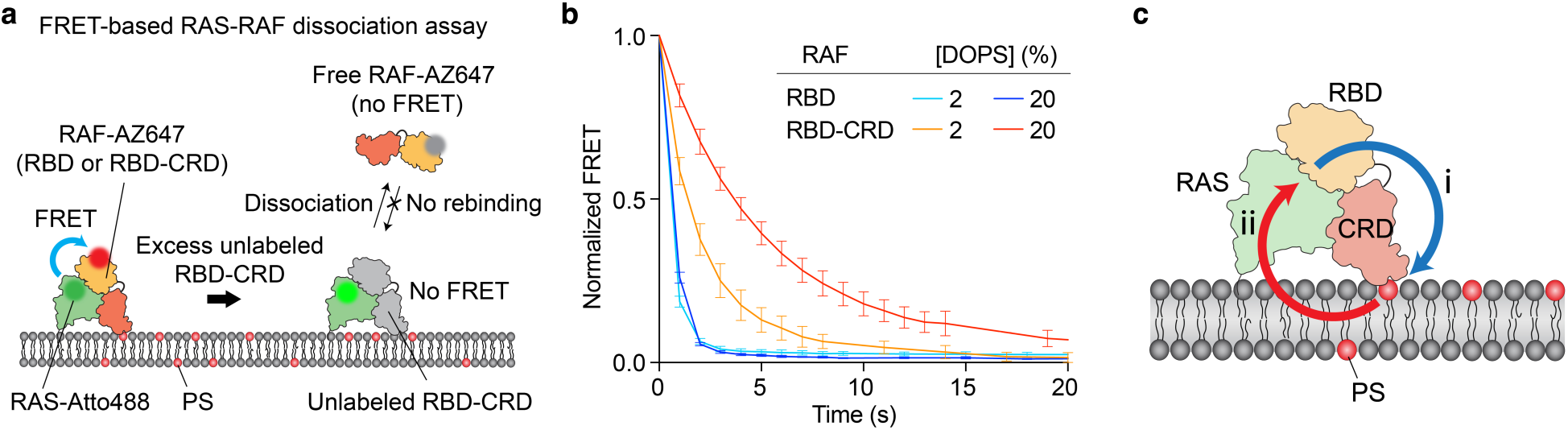
Lipid–CRD engagement enhances the affinity of protein–protein interaction in the RAS:RBD-CRD complex. **a** Overview of the FRET-based competitive assay used to measure RAS-RAF dissociation kinetics. **b** FRET dissociation curves for RBD and RBD-CRD on RAS-functionalized membranes containing 2% or 20% PS (*N*=3). The error bars represent the standard deviations. RAS surface density: ∼500 µm^-2^. Unlabeled RBD-CRD concentration: 3 µM. **c** Schematic illustration of the positive cooperativity between protein–lipid and protein– protein interactions in RAF at the membrane surface. RBD binding to RAS promotes lipid engagement by the CRD (i), which in turn reinforces the association between RAS and RBD– CRD (ii).

### A linker connecting RBD and CRD is crucial for positive cooperativity

We aimed to identify key structural features that mediate the positive cooperativity between RBD and CRD on membranes. X-ray crystallography and solution NMR studies reported that the RBD and CRD simultaneously bind to KRAS through distinct interfaces, forming a trimeric complex^25,47^. In this arrangement, the six-amino-acid-long linker connecting the RBD and CRD plays an essential role in forming the extended structure for KRAS binding by positioning these two tandem domains within close proximity (Fig. 4a). The residues involved in trimer formation are highly conserved across all RAS isoforms^47^. Guided by the KRAS:RBD-CRD(CRAF) structure, we investigated how these interactions mediate the positive cooperativity between the two domains, focusing on regulating membrane binding kinetics. We introduced double alanine mutations at L136 and T178 in the RBD-CRD construct (RBD-CRD^L136A/T178A^) (Fig. 4a). L136, located within the linker region, buries itself within a hydrophobic pocket with F130 in the RBD. T178, located in the CRD, hydrogen-bonds with the C-terminal helix α5 of RAS. These double mutations are expected to disrupt both the linker–RBD and CRD–RAS interactions. Yet, only marginal binding differences were observed between the wild-type and mutant constructs on RAS-functionalized membranes containing 20% PS (red and blue, Fig. 4b), indicating that the CRD has sufficient mobility to change its binding interface with RAS, as previously observed in molecular dynamic (MD) simulations^60^. The linker region contains a rigid proline residue and forms multiple hydrophobic and hydrogen bonds with the RBD and CRD. These collective properties of the linker may be crucial for orienting the CRD to enable effective lipid engagement. Next, we replaced the entire linker sequence with a flexible GGGGGS linker of the same length, which caused a substantial decrease in membrane binding (yellow, Fig. 4b). To examine linker length effects, we attempted to purify a construct featuring an extended linker of four GGGGGS repeats (24 amino acids total), but these efforts were unsuccessful. However, we characterized a construct (4×GGGGGS/T178A) combining the extended linker with the T178A mutation, which showed an even greater reduction in membrane binding (cyan, Fig. 4b). We attribute this decrease primarily to the extended, flexible linker, since T178A had no effect, as shown by the double mutant construct. Notably, the association rate remained largely unchanged across all four constructs (Fig. 4c), while differences in membrane binding affinity were mainly due to changes in the membrane dissociation rate (Fig. 4d). These findings indicate that the linker does not affect the initial association between RBD and RAS; rather, it mediates functional coupling that influences subsequent steps in membrane dissociation, such as CRD– lipid interaction and RBD–RAS unbinding.

**Fig. 4:**
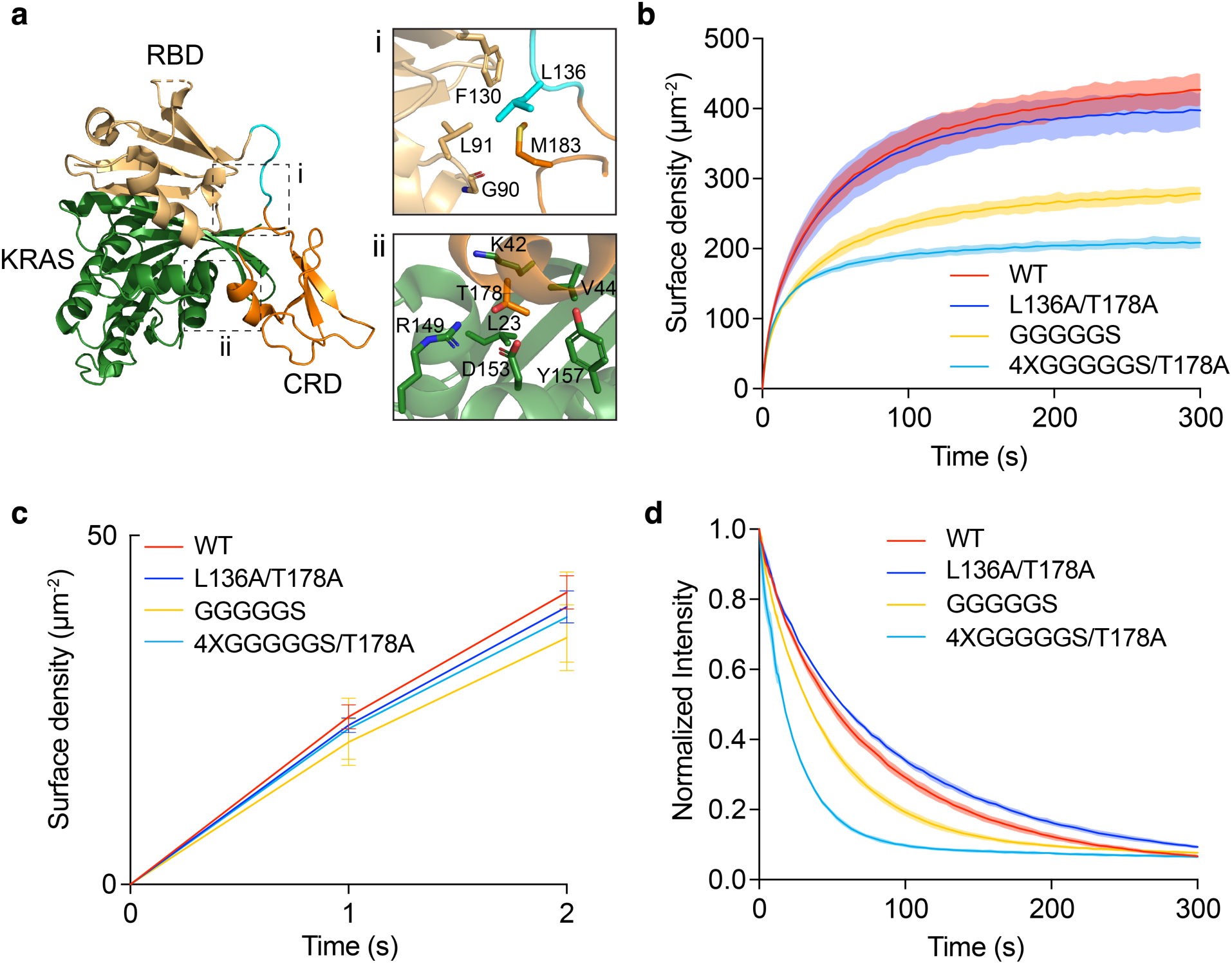
The short linker connecting RBD and CRD is critical for the positive cooperativity. **a** Structure of KRAS:RBD-CRD(CRAF) trimer complex (PBD:6XI7). Inset (i) shows residues interacting with L136, which is part of the interdomain linker, and inset (ii) shows residues interacting with T178 at the CRD and RAS interface. **b** Binding kinetics of WT RBD-CRD and an L136A/T178A mutant that disrupt CRD and RAS interaction, and linker mutants with various linker lengths (*N*=3). The shaded regions represent the standard deviations. **c** Zoom-in of the initial binding curves for WT and mutant RBD-CRD constructs shown in (b). The error bars represent the standard deviations. **d** Ensemble free dissociation measurements of WT and mutant RBD-CRD constructs (*N*=3). The shaded regions represent the standard deviations. All experiments were performed at 10 nM RAF on 20% PS membranes functionalized with RAS at a density of ∼500 µm^-2^.

### RAS density modulates RAF membrane residence through lateral rebinding

We found that the dissociation kinetics of RAF (RBD-CRD) differ markedly between free dissociation assays and competitive FRET assays. In free dissociation assays (red, Fig. 5a), the half-life was about 77 seconds—an order of magnitude longer than the 4-second half-life observed in competitive FRET assays (blue, Fig. 5a). The key difference between these two assays resides in the allowance of rebinding pathways. In FRET assays, rebinding from both solution and the membrane is blocked by excess unlabeled CRAF competing with the labeled CRAF. In contrast, free dissociation assays only prevent solution-based rebinding via continuous buffer flow, leaving membrane-based rebinding possible. This observation suggests that prolonged RAF membrane residence involves a lateral rebinding mechanism, where RAF briefly remains on the membrane via its lipid interactions and can subsequently rebind to another RAS molecule. In cases where RAF fails to rebind to RAS, it rapidly dissociates from the membrane. Based on these results, we propose a kinetic model for RAF membrane dissociation (Fig. 5b). The process begins with RAS unbinding, resulting in a transient RBD-CRD:PS intermediate. This intermediate then unbinds from lipids, leading to complete dissociation of RAF into solution. Alternatively, it can rebind laterally to RAS, restoring the RAS:RBD-CRD:PS complex and extending its membrane residence time. We exclude the scenario where RBD-CRD first loses lipid contact yet remains bound to RAS, as a lipid-free RAS:RBD-CRD complex was not observed in single-particle tracking experiments (Fig. 2e).

**Fig. 5:**
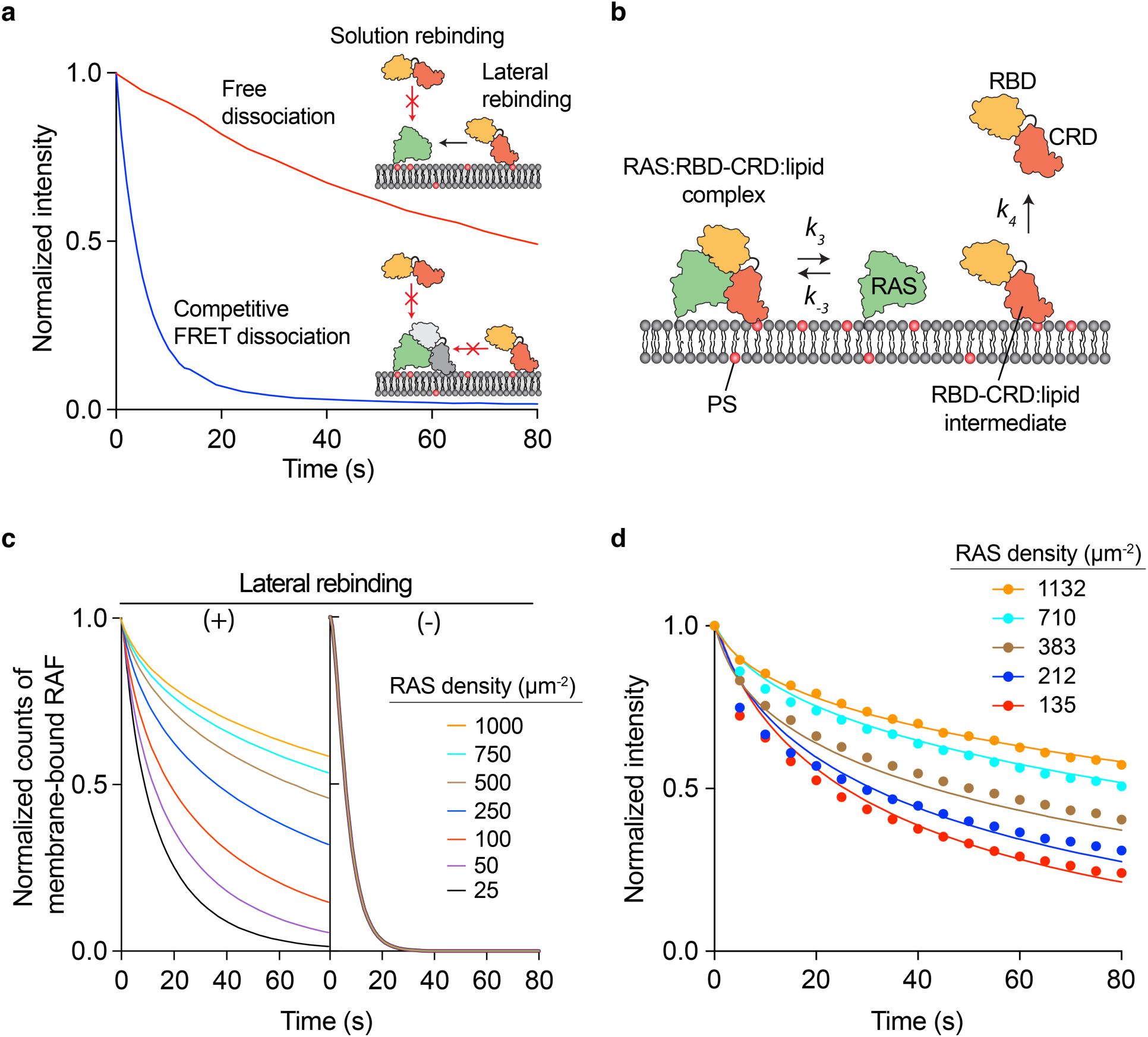
RAS surface density modulates membrane residence time of RAF via lateral rebinding. **a** Comparison between free dissociation (Fig. 2c) and competitive FRET dissociation (Fig. 3b) curves for RBD-CRD on RAS-functionalized membranes containing 20% PS. **b** Schematic illustration of RAF RBD-CRD dissociation mechanism involving lateral rebinding to RAS. **c** Simulations for RAF RBD-CRD membrane dissociation kinetics at various RAS densities, with (+) and without a rebinding pathway (−). **d** Experimental free dissociation kinetics of RBD-CRD under different RAS densities on 20% PS membranes. RBD-CRD concentration: 20 nM. (See Supplementary Information for detailed methods for kinetic simulation and the estimated kinetic constants).

Kinetic simulations further illuminate how RAS nanoclusters regulate RAF membrane dissociation kinetics. The lateral rebinding of a transient RBD-CRD:PS intermediate to RAS is a second-order process whose rate is linearly proportional to RAS surface density. Consequently, higher RAS densities increase RAF dwell times on membranes (left, Fig. 5c). In contrast, without lateral rebinding, dissociation follows a two-step, irreversible first-order pathway, removing any dependence on RAS density (right, Fig. 5c). To validate the lateral rebinding mechanism, we measured RAF dissociation kinetics at various RAS densities, revealing a strong modulation of membrane dissociation by RAS density. In particular, RBD-CRD remained on the membrane significantly longer at higher RAS densities (Fig. 5d). Although more complex models could be constructed, this two-step dissociation model successfully and quantitatively accounts for TIRF desorption data. The calculations and estimated rate constants for RAS unbinding (*k*_3_), RAS rebinding (*k*_–3_), and lipid unbinding (*k*_4_) are presented in Supplementary Table 2 and its associated text. The rebinding mechanism may play a pivotal role in RAF activation, particularly in the context of RAS nanoclusters known to modulate MAPK signaling outputs. Overall, this mechanism provides a kinetic framework for understanding how RAS nanoclusters function as signaling hubs, regulating RAF membrane residence time and activity.

## Discussion

An important unresolved question in RAF signaling concerns the mechanisms that trigger structural rearrangements to release autoinhibition and facilitate the completion of its multistep activation process. Since RAF activation occurs exclusively on the plasma membrane, understanding the mechanisms that modulate membrane binding kinetics is essential. Yet, our current knowledge remains largely qualitative, lacking detailed mechanistic explanations.

The present study provides new insights into RAF activation by providing a quantitative kinetic model of membrane interaction and uncovering the mechanism by which the tandem RBD and CRD cooperate to generate emergent properties that facilitate activation. Using a combination of TIRF microscopy, FCS, and SLB technology, we demonstrated that the RBD and CRD of CRAF synergistically enhance membrane binding affinity. While the affinity between CRD and PS lipids is weak—nearly undetectable at physiological RAF concentrations, it induces a profound effect when arranged in tandem with RBD, resulting in approximately a 30-fold increase in membrane binding affinity compared to the isolated RBD. The kinetic analysis revealed that the enhanced affinity is solely due to an extended membrane dwell time, i.e., a decrease in membrane dissociation rate, *k*_off_. The CRD and PS lipid interaction shows no measurable impact on *k*_on_, ruling out a major role of CRD in the initial membrane association step of CRAF. Instead, the RBD and RAS interaction strictly dictates the membrane association rate, *k*_on_. These findings support the conclusion that CRAF membrane association occurs in two kinetically distinct steps: first, RBD binding to RAS, followed by CRD engagement with lipids.

The two-step association mechanism is compatible with the fully autoinhibited structure observed in BRAF, in which the RBD remains exposed for RAS binding^40,42^. Such an association mechanism will enable RAF to exploit abundant PS lipids as a specific cue in response to RAS activation.

The tandem arrangement of the RBD and CRD, connected by a six-amino-acid linker, raises the question of whether these two domains influence each other’s functions. Our study demonstrates that positive cooperativity emerges specifically within membrane environments, significantly slowing membrane dissociation kinetics. We identified three interwoven mechanisms contributing to this behavior. First, RBD binding to RAS promotes the lipid-sensing ability of the CRD, thereby enhancing overall membrane binding. This cooperative interaction provides the mechanistic basis for two-step association, with the flexibility and length of the short linker acting as critical determinants. Second, membrane stabilization by the CRD arises from a synergistic effect, in which CRD–lipid interaction, enabled by RBD–RAS binding, in turn reinforces RBD–RAS association. This finding aligns with computational studies, which predict that CRD–membrane interactions reduce fluctuations in the RAS:RBD complex, thereby enhancing its stability^59^. Lastly, RAF dissociation involves a lateral rebinding mechanism.

Following RAS unbinding from the RAS:RBD-CRD:PS complex, a transient RBD-CRD:PS intermediate can remain on the membrane and laterally rebind to RAS, significantly extending membrane residence at high RAS density. In this mechanism, a weak CRD–lipid interaction plays an essential role in momentarily holding the intermediate on the membrane. Previous computational and experimental studies have shown that RBD induces local anionic lipid enrichment independently of RAS, thereby enhancing the overall membrane affinity of RBD-CRD^61^. Consistent with this, we observed stronger binding of RBD-CRD on RAS-free membranes. The RBD-induced local enrichment of anionic lipids may support lateral rebinding by slowing the dissociation of the transient RAS-free RBD-CRD:PS intermediate. Moreover, the orientational flexibility of the RBD-CRD, observed in MD simulations and NMR studies, likely facilitates lateral rebinding by enabling efficient sampling of conformational ensembles compatible with RAS binding^25,61^. Overall, synergistic enhancement of the RBD and CRD affinities for their respective binding partners, combined with lateral rebinding, regulates RAF membrane-binding kinetics. Coexistence of protein and lipid binding is broadly observed in many signaling proteins^62^. Lateral rebinding may be a common mechanism to extend membrane residence via the multivalency of reversible weak interactions.

Our study revealed that lateral rebinding has a critical functional outcome: it prolongs RAF membrane residence at higher RAS·GTP densities. This raises two important questions: how does membrane dwell time modulation facilitate RAF activation, and what are the functional benefits of this mechanism? This RAS density-dependent change in membrane residence is biologically significant, given that RAF activation occurs via RAS nanoclusters^43–45^. RAS nanoclusters function as an analog–digital–analog converter, linearly transducing continuous epidermal growth factor (EGF) input into digital, fully active clusters^44,45^. A key behavior is that as RAS×GTP concentration increases, the number of RAS nanoclusters rises proportionally while the cluster size remains quantized^44^. Because they rely on collective properties, nanoclusters are inherently less sensitive to fluctuations in activating inputs and intrinsic biochemical noise, thus providing robust, high-fidelity transmission^43^. However, only about 40% of RAS forms nanoclusters, leaving the majority in a monomeric state that is more vulnerable to cellular fluctuations^43,44^. This observation suggests that additional mechanisms are required to favor RAF activation at RAS nanoclusters over monomeric RAS, thereby achieving the high-fidelity signaling observed. The current model emphasizes a local concentration effect, wherein RAS nanoclusters enrich RAF to promote the formation of active dimers^63^. Here, we propose an additional complementary mechanism in which RAF discriminates between monomeric and clustered RAS based on membrane residence time. We hypothesize that RAF activation is regulated by a type of kinetic proofreading mechanism^64,65^, whereby molecules that remain on the membrane for extended periods—such as those within RAS nanoclusters—are disproportionately more likely to become activated. RAF possesses two necessary features for a cytosolic enzyme to achieve kinetic proofreading on membranes: (i) elongation of dwell time, and (ii) a multistep activation process^64^. Under this mechanism, enzymes with slow dissociation kinetics that dwell on membranes for sufficiently long periods can complete the multistep activation process, whereas short-dwelling species dissociate prematurely before activation is complete, as previously demonstrated in the guanine nucleotide exchange factor SOS^66^.

Based on previous studies and our quantitative kinetic analysis, we propose a model for RAF activation on membranes (Fig. 6). In the resting state, RAF exists as an autoinhibited monomer complexed with MEK and a 14-3-3 dimer in the cytosol^39^. Activation begins when RAS·GTP recruits RAF via binding to its accessible RBD (Step i in Fig.6)^40,42^. During this initial interaction, the CRD is positioned near the membrane by a short six-amino-acid-long linker, thereby promoting lipid engagement (Step ii in Fig. 6). This step requires CRD extraction, potentially driven by conformational fluctuations or destabilization of RBD and 14-3-3 contacts^40,42^, although the precise mechanism remains to be determined. Successful lipid engagement by the CRD strengthens the RBD–RAS association and stabilizes an open RAF conformation. Unbinding from RAS yields a RAF:PS intermediate, which can be restored to a RAS:RAF:PS complex via lateral rebinding to RAS (Step iii in Fig. 6). This rebinding process— particularly efficient within RAS nanoclusters—further prolongs RAF membrane residence. In our model, extended dwell time is achieved through a series of transient RAS interactions within nanoclusters, dynamically replenished from the monomeric pool. If a RAF:PS intermediate fails to rebind to RAS, it dissociates into the cytosol before reaching full activation. (Step iv in Fig. 6) Persistent CRD–lipid engagement sustains displacement of 14-3-3 from the CR2 region, enabling dephosphorylation by the SHOC2–MRAS–PP1 complex and preventing reversion to the autoinhibited state. The open RAF monomer then forms active dimers, stabilized by rearranged 14-3-3 dimers binding to C-terminal phosphoserine (Step v in Fig. 6). Along with local concentration effects, extended membrane residence via lateral RAS rebinding supports kinetic proofreading within RAS nanoclusters to complete this multistep activation process—encompassing global domain rearrangements, SHOC2–MRAS–PP1C-mediated dephosphorylation, dimer formation, and *trans* auto-phosphorylation. The extended dwell time also likely prolongs the lifetime of active RAF, allowing it to perform more catalytic reactions. Conversely, sparse RAS yields insufficient residence to complete activation, thereby averting spontaneous signaling. Given that dwell time modulation is a recurring theme in multistep-activation enzymes^66–68^, kinetic proofreading may serve a regulatory mechanism in many membrane-proximal signaling reactions.

**Fig. 6:**
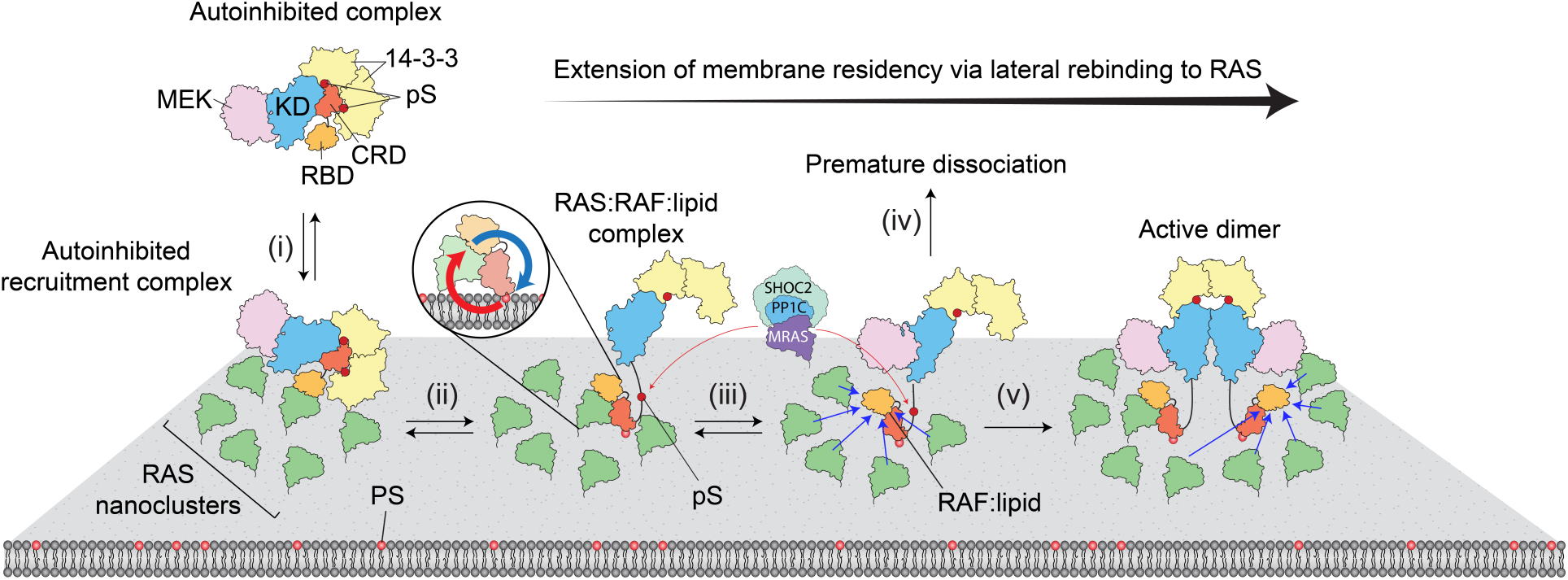
A model for membrane-dependent activation of RAF. RBD binding to RAS initiates RAF membrane recruitment (i). RAS engagement within the membrane environment displaces 14-3-3 dimers, exposing the CRD, which then engages lipids and transitions into an open conformation (ii). Successful engagement of both RBD–RAS and CRD–lipid interactions mutually enhances their affinities (depicted in the inset), further stabilizing RAF on the membrane. High-density RAS nanoclusters prolong RAF membrane residence through lateral rebinding (iii). Failure to rebind leads to RAF dissociation before full activation (iv). During the extended membrane residence, the SHOC2-MRAS-PP1C complex dephosphorylates the phosphoserine residue in the CR2 region (pS365 for BRAF, pS259 for CRAF), preventing reversion to a closed conformation. Ultimately, RAF completes multistep activation by forming an active dimer (v).

Our in vitro reconstitution system using supported lipid bilayers has successfully quantified the cooperativity between the RAF RBD and CRD, providing a promising foundation for further study. We plan to extend this approach by incorporating more complex lipid compositions and KRAS with its farnesyl modification to better mimic native nanoclusters^57^.

Additionally, integrating the autoinhibited full-length RAF:14-3-3:MEK complex with the SHOC2-MRAS-PP1C module will allow us to examine how membrane binding kinetics influence activation outcomes under physiologically relevant conditions. These advancements will enable us to rigorously recapitulate the sequential steps of RAF activation and elucidate the underlying molecular mechanisms with high resolution.

## Methods

### Plasmid cloning

CRAF constructs were cloned into modified 2CT pET plasmids containing an N-terminal TwinStepII tag, maltose-binding protein (MBP), and a TEV protease cleavage site, followed by the protein of interest (2CT pET-TwinStrep-MBP). For mNeonGreen labeling, three repeats of the GGGGS amino acid linker were appended to the C-terminus, followed by the mNeonGreen tag. For sortase-mediated labeling with Gly-Gly-Gly-AZ647 (Vector Laboratories), a C-terminal LPETGG sequence was introduced. CRAF domains (51-131 for RBD, 51-188 for RBD-CRD, and 138-188 for CRD) were PCR-amplified and assembled using Gibson Assembly to yield TwinStrep-MBP-TEV-CRAF-mNG. Additionally, HRAS(1-184, C118S) and the SOS1 catalytic domain (SOS_cat_, 565-1049) were cloned into a pProEXHTB plasmid to construct His_6_-TEV-HRAS and His_6_-TEV-SOS_cat_, respectively.

### Protein Purification

CRAF constructs were expressed in Rosetta(DE3) cells. An overnight preculture was grown, then diluted into 1 L, and further grown at 37 °C. Protein expression was induced with 30 µM IPTG at an O.D. of 0.6. Cells were harvested by centrifugation after five hours of induction, frozen, and stored at −80 °C until purification. For purification, cell pellets were resuspended in lysis buffer consisting of Buffer A (20 mM HEPES [pH 7.3], 500 mM NaCl, 1 mM tris(2-carboxyethyl) phosphine [TCEP], and 10% glycerol) supplemented with 15 µg×mL^-1^ DNase and 1 mM phenylmethylsulfonyl fluoride (PMSF). Cells were lysed, and the lysate was clarified by centrifugation before purification using fast protein liquid chromatography (FPLC) on an NGC system (Bio-Rad). An initial capture step was performed using a Strep-Tactin affinity column (Strep-TactinXT 4Flow, IBA Lifesciences), followed by elution with buffer A supplemented with 50 mM biotin. Selected fractions were dialyzed overnight with TEV protease at 4 °C. The cleaved tag and uncleaved protein were removed by reintroducing the cleaved sample onto a Strep-Tactin column. Further purification was performed using size-exclusion chromatography (Superdex 75 Increase HiScale, Cytiva). Collected fractions were analyzed by SDS-PAGE, pooled as appropriate, aliquoted, snap-frozen in liquid nitrogen, and stored at −80°C. HRAS, SOS, and NF1 were expressed and purified as described elsewhere^49^.

### SLB preparation

Glass coverslips (D263 Schott glass, Ibidi) were first cleaned by sonication in a 1:1 isopropanol/water mixture for 15 minutes, followed by incubation in 2% warm Hellmanex III for 30 min. The substrates were then etched for 10 minutes in a piranha solution (3:1 H₂SO₄/H₂O₂). The coverslips were assembled on six-channel flow chambers (sticky-Slide VI 0.4, Ibidi). SLBs were formed by fusion of small unilamellar vesicles (SUVs) as described elsewhere^49^. For SLBs containing 2% PS, the lipids were mixed at 97 mol% 1,2-dioleoyl-sn-glycero-3-phosphocholine (DOPC), 2 mol% 1,2-dioleoyl-sn-glycero-3-phospho-L-serine (DOPS), and 1 mol% 1,2-dioleoyl-sn-glycero-3-phosphoethanolamine-N-[4-(p-maleimidomethyl) cyclohexane-carboxamide] (MCC-DOPE). For SLBs containing 20% PS, the lipids were mixed at 77 mol% DOPC, 20 mol% DOPS, and 3 mol% MCC-DOPE. RAS was functionalized via a covalent reaction between the terminal cysteine residues and the maleimide group of MCC lipids as described elsewhere^49^. Typically, a concentration of 0.1–0.5 mg·mL^-1^ RAS in PBS was incubated for 2.5 hours. The reaction was terminated by incubation of 10 mM beta-mercaptoethanol (BME) for 10 minutes. Nucleotides bound to RAS were stripped with 10 mM EDTA in HBS (40 mM HEPES, 150 mM NaCl, pH 7.4). Finally, RAS was loaded with the desired nucleotide by overnight incubation of the sample with either 1 µM Atto488-GDP (Jena Bioscience) or 10 µM GppNp in HBS supplemented with 5 mM MgCl_2_ (hereafter referred to as HBS/MgCl_2_). For mNG FCS calibration measurements, SLB containing 96 mol% DOPC and 4 mol% 1,2-dioleoyl-sn-glycero-3-[(N-(5-amino-1-carboxypentyl)iminodiacetic acid)succinyl] (DOGS-NTA) were prepared by vesicle fusion method identical to SLBs containing PS lipids. A concentration ranging from 1-10 nM of His_10_-mNG was incubated in PBS for 10 minutes to yield various surface densities. The unbound mNG was rinsed with PBS.

### Microscopy

Imaging was performed using a Nikon Ti-2 microscope equipped with an Apo TIRF 100× oil immersion objective and an EMCCD camera (iXon Life 897, Oxford Instruments). For RAS density and RAF-mNeonGreen measurements, a 488 nm laser (Coherent), filtered through a laser cleanup filter (ZET488/10x, Chroma Inc.), was directed onto the sample using a dichroic mirror (zt488/640rpc, Chroma Inc.). The fluorescence was collected through two stacked emission filters (ZET488/640m and ET535/70m, Chroma Inc.). RAF-AZ647 was excited with a 640 nm laser (Coherent), filtered through a laser cleanup filter (LD01-640/8, Semrock) and directed by the same dichroic mirror (zt488/640rpc). The fluorescence was collected through two stacked emission filters (ZET488/640m and ET706/95m, Chroma Inc.).

### Ensemble adsorption and desorption

In ensemble experiments, SLBs were initially prepared with RAS loaded with the fluorescent nucleotide Atto488-GDP to determine surface density. Nucleotide exchange to GTP was facilitated by incubation of 100 nM catalytic domain of SOS and 100 µM GTP in HBS/MgCl_2_ for 6 min. Active RAS·GTP-functionalized membranes are nonfluorescent and suitable for subsequent binding assays with RAF fused with mNeonGreen. The binding measurements were performed in HBS/MgCl_2_ supplemented with 0.1 mg·mL^-1^ casein, 1 µM GTP, and 1 mM β-mercaptoethanol (BME) (hereafter referred to as CGB buffer). Upon reaching a plateau in the binding curve (typically in 5 min), desorption measurements were conducted. Desorption was initiated by injecting 200 µL of CGB buffer, followed by continuous buffer flow at a rate of 1 mL·min^-1^ using a syringe pump. Background signals, *I*_bg_, were obtained from the average intensities of the frames prior to RAF injection and were subtracted from the raw binding intensities. The intensity values were then divided by the maturation efficiency, *M*_Eff_, of RAF-mNeonGreen proteins. The solution contribution, *I*_soln_, of mNeonGreen was then subtracted to calculate the binding/desorption net intensities, *I*, for analysis.

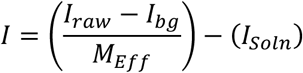

The net intensity was converted to the surface density using the TIRF-FCS calibration curve.

### Competitive FRET Dissociation

Competitive desorption assays were performed using Förster Resonance Energy Transfer (FRET) to quantitatively assess the stability of the RAS:RAF complex. The donor Atto488-GppNp (Jena Bioscience) was loaded on RAS (RAS-Atto488-GppNp) and the acceptor AZ647(Vector Laboratories) was conjugated to either RBD or RBD-CRD of CRAF (RAF-AZ647). The RAS:RAF complex was formed by incubating 20 nM RAF-AZ647 with RAS tethered on SLBs for 5 min. 3 μM unlabeled RBD-CRD was subsequently introduced. FRET measurements were acquired and analyzed as described elsewhere^69^. Three fluorescence signals were recorded: donor/acceptor (DA), acceptor/acceptor (AA), and donor/donor (DD). DD and AA are regular fluorescence signal acquired with the optical configurations described in the Microscopy section. The DA signal was acquired with a 488 nm laser excitation and emission configured for 640 nm channel. The measured DA signal, representing total acceptor fluorescence under donor excitation, is described by the following equation:

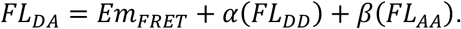

Here, *FL*_DA_ is the total acceptor emission under donor excitation, *Em*_FRET_ is the sensitized emission due to FRET, α(*FL*_DD_) accounts for donor emission bleed-through into the acceptor channel, and β(*FL*_AA_) accounts for direct acceptor excitation at the donor excitation wavelength. The FRET signal was calculated by subtracting contribution of α(*FL*_DD_) and β(*FL*_AA_) from *FL*_DA_.

### Single-molecule recruitment

Single-particle recruitment measurements of RAF-AZ647 were conducted in CGB buffer on RAS·GTP-functionalized membranes. TIRF measurements were acquired using a 20 ms exposure time in stream mode, with the 640 nm laser power set to 5 mW. Single-molecule trajectories were detected and built using the TrackMate plugin in ImageJ^70^. Cumulative binding events were then counted and plotted over the duration of the measurement using custom MATLAB scripts. The slope of this plot, normalized by RAS density, was used to determine *k*_on_^49^.

### Single-molecule tracking

Single-molecule diffusion experiments were conducted using an oxygen scavenging buffer consisting of CGB buffer supplemented with 0.32 mg·mL^-1^ glucose oxidase from Aspergillus niger (Serva), 0.05 mg·mL^-1^ catalase from bovine (Sigma-Aldrich), 2 mM Trolox (Sigma-Aldrich), 1 µM GTP, and 20 mM glucose. RAF-AZ647 and RAS-Alexa647-GppNp diffusion measurements were performed on a RAS·GTP-functionalized SLBs. Single-molecule trajectories were detected and built using the TrackMate plugin in ImageJ^70^. The step size distribution was calculated using custom MATLAB scripts. A Brownian diffusion model, described below, was used to fit the step size distribution of each measurement:

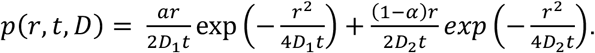

Diffusion coefficients, *D_1_* and *D_2_*, and relative population, for the fast species, *α* were calculated from the corresponding fitting. Step size is represented by *r*, at an experimental time interval, *t*.

### Fluorescent Correlation Spectroscopy

Measurements were performed on a custom-built FCS setup integrated with a Nikon Ti-2 microscope. A SuperK Evo HP white-light laser source (NKT Photonics) was reflected by a dichroic mirror (zt488rdc, Chroma Inc.) and filtered through a cleanup filter (LL01-488, Semrock) to select a 488 nm wavelength. The beam was coupled into a polarization-maintaining single mode fiber (PM-S405-XP, Thorlabs) via a fiber coupler (PAF2-A4A, Thorlabs). The beam collimated via a reflective collimator (RC08FC-P01, Thorlabs) was directed to the back port of the microscope and then to the objective by a quad-band dichroic filter cube (ZT405/488/561/640rpc, Chroma Inc.). The excitation beam was focused by Apo TIRF 100× oil immersion objective. Fluorescence emission from the sample was spatially filtered through a 50 µm pinhole, collimated by an achromatic doublet lens (AC254-125-A, Thorlabs), reflected by a dichroic mirror (550lpxr, Chroma Inc.), and further filtered through a bandpass filter (ET520/40m, Chroma Inc.). It was then refocused by another achromatic doublet lens (AC254-75-A, Thorlabs) onto an avalanche photodiode (SPCM-AQRH-16, Excelitas Technologies) connected to a time-correlated single-photon counting module (PicoHarp 300, PicoQuant). Data acquisition was performed using PicoHarp software (PicoQuant) and analyzed in MATLAB and Prism (GraphPad). The autocorrelation curve, *G*(τ), was fitted to a two-dimensional Gaussian diffusion equation to calculate surface density of RAF and RAS.

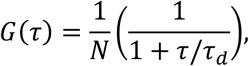

where *N* is the number of particles within the focal area, τ is the time delay, and τ_9_ is the correlation time. The focal radius was determined to be 0.169 µm and was used to calculate the protein density.

## Data availability

The authors declare that the data supporting the findings of this study are available within the paper and its Supplementary Information files or from the corresponding author upon request.

## Acknowledgments

This study was supported by CAREER Award MCB-2145852 to Y. L. A. J. S. was supported in part as a Fellow of the Rees-Stealy Research Foundation.

## Author contributions

A.J.S. expressed and purified all proteins, with assistance from K.T. and A.C. A.J.S., K.T. and Y.K.L. performed and analyzed ensemble and single-molecule binding assays, with assistance from J.G, A.C, and A.M. J.G. performed FCS experiments. Y.K.L. designed the experiments and directed the project. A.J.S. and Y.K.L. drafted the manuscript with input from all authors.

## Competing interests

The authors declare no competing interests.

Correspondence and requests for materials should be addressed to Youngkwang Lee.

## Notes

### Competing Interest Statement

The authors have declared no competing interest.

