## Supplementary Information for "Positive cooperativity between RAS-binding and cysteine-rich domains regulates RAF membrane binding kinetics via lateral rebinding"

**Supplementary Information**
Supplementary Figures 1-10
Supplementary Table 1
Supplementary Text: Estimating kinetic rate constants involved in RAF membrane dissociation

### 1. Supplementary figures

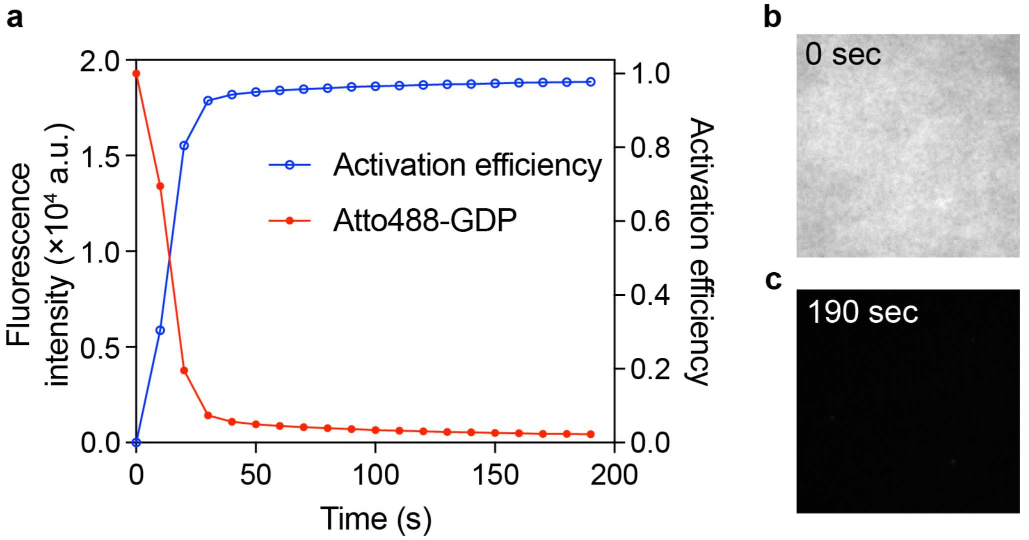


**Supplementary Figure 1. RAS activation via GTP nucleotide exchange using the SOS catalytic domain. a** Fluorescence intensity of Atto488-GDP-bound RAS decreases over time as SOS replaces Atto488-GDP with unlabeled GTP. The activation efficiency increases inversely to the fluorescence drop and is calculated by $|I_o - I(t)|/I_0$. **b,c** Fluorescence microscopy images of Atto488-GDP-bound RAS before (b) and after activation (c). Near-complete RAS activation is achieved within three minutes of SOS-mediated nucleotide exchange (SOS concentration: 100 nM). RAS density on the membrane is ~1200 μm$^{-2}$, and the membrane composition is 77% DOPC, 20% DOPS, and 3% MCC-DOPE.

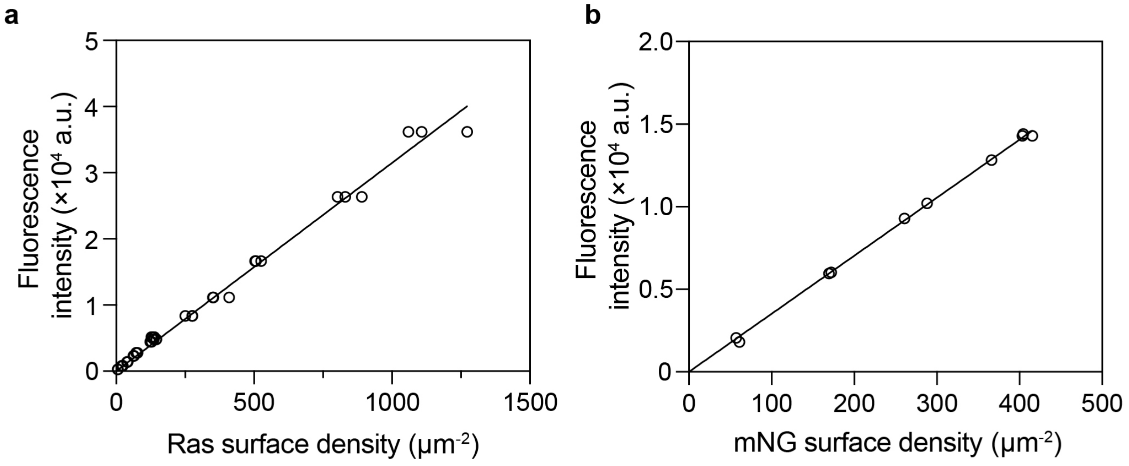


**Supplementary Figure. 2. TIRF calibration.** The surface density of protein on supported membranes was determined by TIRF intensity, calibrated against FCS density measurements. **a** TIRF calibration of Atto488-GDP-labeled RAS**. b** TIRF calibration of His6-mNeonGreen (mNG) used for RAF-mNG constructs. The membrane composition is 77% DOPC, 20% DOPS and 3% MCC-DOPE for RAS calibration, and 96% DOPC and 4% NDG-NTA-DOPE for mNG calibration.

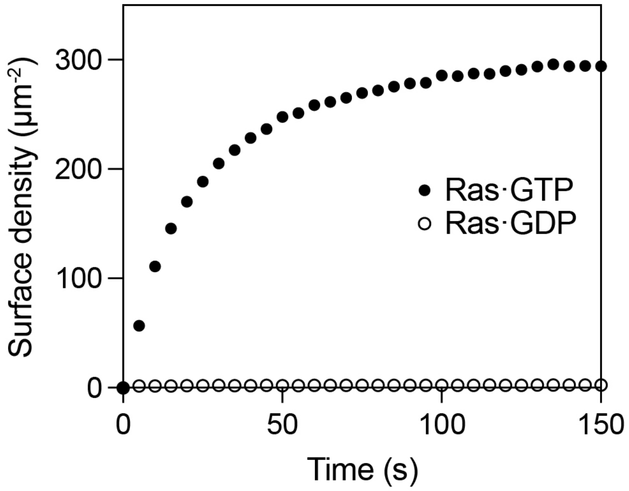


**Supplementary Figure 3. RBD-CRD binding on active and inactive RAS-functionalized membranes.** Shown are the binding kinetics of 20 nM RBD-CRD on supported membranes functionalized with RAS·GTP (~240 $\mu m^{-2}$) and RAS·GDP (~450 $\mu m^{-2}$). RBD-CRD displays robust binding to RAS·GTP membranes, whereas negligible binding is observed on RAS·GDP membranes. The membrane composition is 77% DOPC, 20% DOPS and 3% MCC-DOPE.

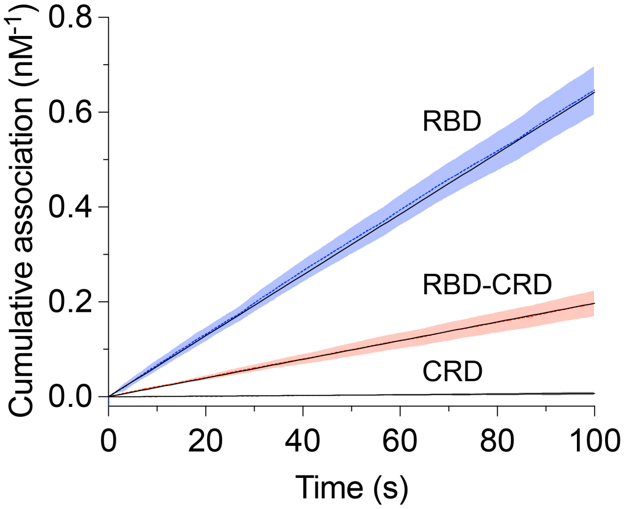


**Supplementary Figure 4. Cumulative RAF binding on RAS-functionalized membranes.** A representative plot of cumulative RAF binding events over time shows a linear increase, indicating a constant rate of binding. The apparent $k_{on}$ value is determined from the slope of this linear fit (black solid line), normalized by RAS density. Shaded bands indicate the standard deviation ($N$=3). The membrane composition is 77% DOPC, 20% DOPS and 3% MCC-DOPE.

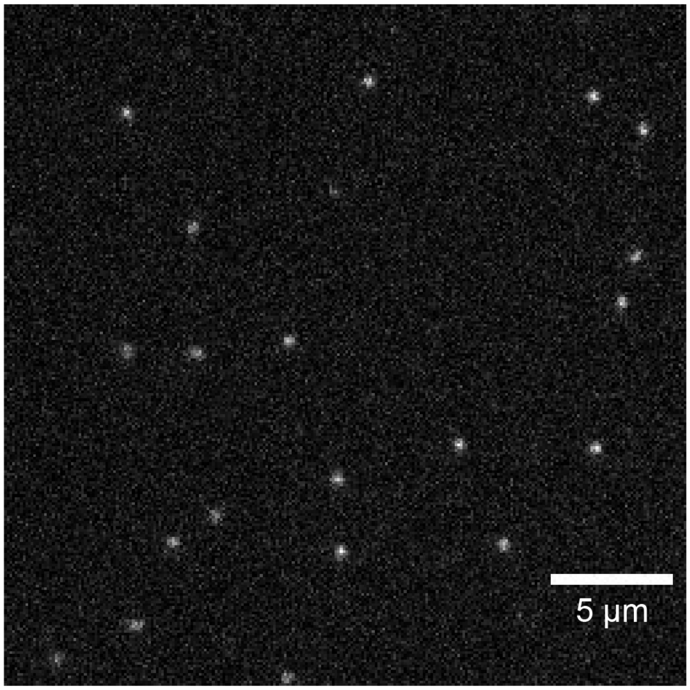


**Supplementary Figure 5. Representative snapshot image showing Alexa647-GppNp-labeld RAS.** For single particle tracking of RAS, a small population of RAS was labeled with Alexa647-GppNp, ensuring a well-dispersed single-molecule density (~20-30 particles per region of interest). Total Ras density: ~550 µm$^{-2}$.

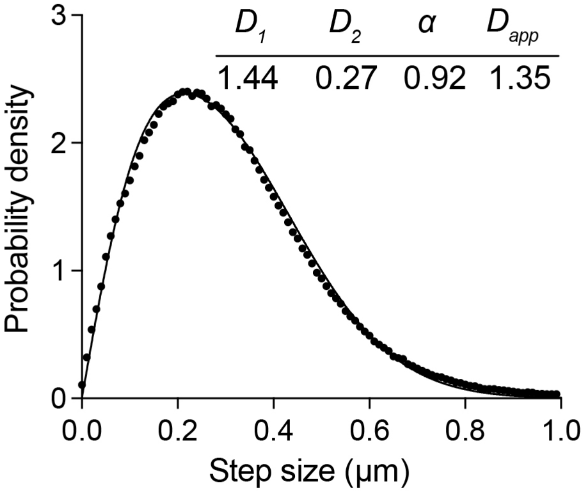


**Supplementary Fig. 6 Step size distribution of Atto565-DOPE lipids on a supported lipid bilayer.** The lipids exhibit two distinct diffusion populations, with the majority species (>90%) diffusing at 1.44 µm²/s. The membrane composition is 80 mol% DOPC, 20 mol% DOPS, and 1x10$^{-5}$ mol% Atto565–DOPE. Images were recorded with a time interval of 10 ms.

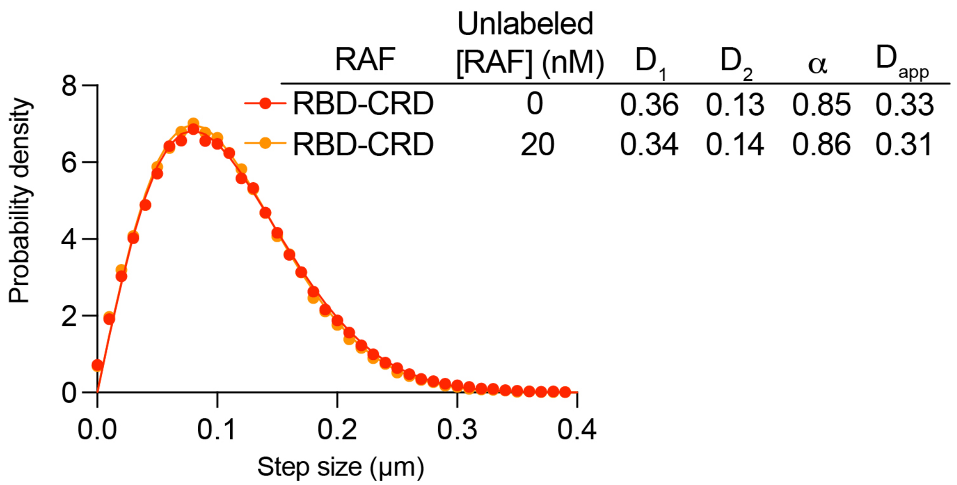


**Supplementary Fig. 7 Diffusion analysis of RBD-CRD in the absence and presence of excess unlabeled Raf.** Diffusion of RBD-CRD remain unchanged at a saturating concentration of unlabeled RBD-CRD (20 nM), confirming that the slower diffusion observed relative to RBD in Fig. 2e is due to lipid interactions with individual RAS:RBD-CRD complexes rather than collective behaviors such as clustering. The experiments were performed on membranes composed of 77% DOPC, 20% DOPS and 3% MCC-DOPE, functionalized with RAS at a density of ~400 $\mu m^{-2}$. 10 pM and 100 pM of RBD-CRD-AZ647 were used for experiments without and with unlabeled RBD-CRD. The images were acquired with a 10 ms time interval.

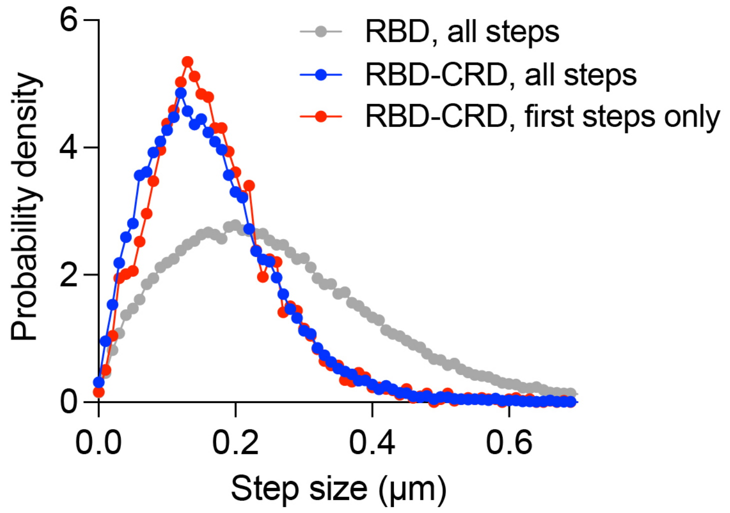


**Supplementary Fig. 8 Distribution of RAF at different time ranges.** RBD-CRD displays similar diffusion during the initial steps of trajectories (first steps) and across the entire trajectories (all steps). The experiments were performed on membranes composed of 77% DOPC, 20% DOPS and 3% MCC-DOPE. RAS density is ~350 $\mu m^{-2}$ for both RBD-CRD and RBD measurements. To minimize blinking effects, the experiments were conducted under fast bleaching conditions in the absence of glucose oxidase/catalase imaging buffer, with 1 mM BME added to eliminate potential oxidative photo effects. Images were acquired with a 20 ms time interval.

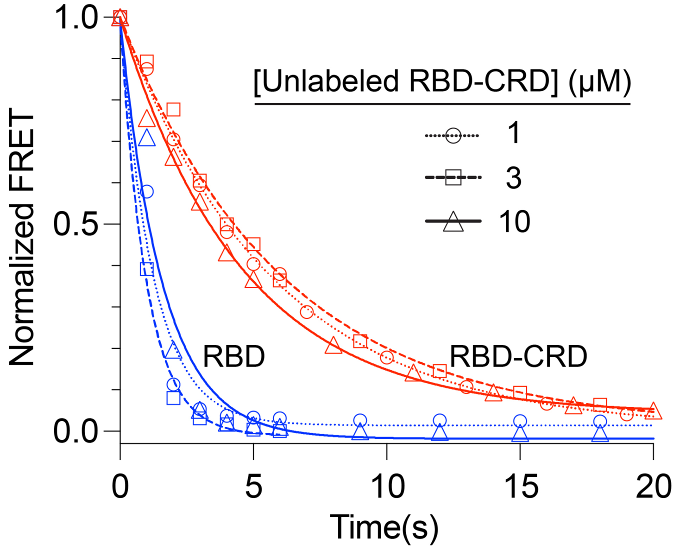


**Supplementary Fig. 9 Competitive FRET dissociation kinetics for RAF-AZ647 constructs with different concentrations of unlabeled RBD-CRD.** Competitive FRET dissociation experiments between RAF-AZ647 and Atto488-GppNp-RAS were performed in the presence of unlabeled RBD-CRD at concentrations ranging from 1 to 10 $\mu$M. All tested micromolar concentrations of unlabeled RBD-CRD are equally efficient at sequestering free RAS upon dissociation of RAF-AZ647 (RBD in blue; RBD-CRD in red). The experiments were conducted on membranes composed of 77% DOPC, 20% DOPS and 3% MCC-DOPE. Ras density is ~400 $\mu m^{-2}$ for RBD and RBD-CRD measurements.

### 2. Estimating kinetic rate constants involved in RAF membrane dissociation

Based on our experimental findings, we propose the following reaction scheme to describe how the CRAF RBD-CRD associate with the membrane:

| Step | Reaction | Rate constants |
|---|---|---|
| **(1)** | RAS(m) + RBD-CRD(s) ↔ RAS:RBD-CRD(m) | $k_1/k_{-1}$ fwd/rev rate constants |
| **(2)** | RAS:RBD-CRD(m) + PS(m) → RAS:RBD-CRD:PS(m) | $k_2$ fwd rate constant |
| **(3)** | RAS:RBD-CRD:PS(m) ↔ RAS(m) + RBD-CRD:PS(m) | $k_3/k_{-3}$ fwd/rev rate constants |
| **(4)** | RBD-CRD:PS(m) → RBD-CRD(s) + PS(m) | $k_4$ fwd rate constant |

(m) and (s) denote the membrane and solution phases, respectively.

The initial step involves RBD binding to membrane-bound RAS, forming RAS:RBD-CRD on the membrane (1). Once the RBD-CRD is membrane-associated via RBD–RAS interaction, the CRD engages lipids (e.g., PS) to form RAS:RBD-CRD:PS (2). We did not observe the reverse reaction of step (2) in our experimental conditions. Dissociation begins with RAS unbinding, creating a transient RBD-CRD:PS intermediate that can laterally rebind to RAS (3) or proceed to the final unbinding step. In the final step, the CRD detaches from the membrane lipid PS (4). The reverse reaction of step (4) was negligible at the concentration used in dissociation assays (20 nM). The stoichiometry between CRD and PS is not well-defined because its promiscuous nature that relies on multiple interdigitated positively charged and hydrophobic residues. For simplicity, we therefore treated this interaction effectively monovalent, representing the convoluted multivalent binding with an apparent rate constant. Overall, the model describes a two-step association (steps 1 and 2) followed by two sequential dissociation steps (steps 3 and 4).

For the free dissociation experiments, only steps (3) and (4) of our reaction scheme are relevant. The corresponding rate laws for these steps are:

$$d[RAS]/dt = k_3[RAS{:}RBD\text{-}CRD{:}PS] - k_{-3}[RAS][RBD\text{-}CRD{:}PS]$$
$$d[RBD\text{-}CRD]/dt = k_4[RBD\text{-}CRD{:}PS]$$
$$d[PS]/dt = k_4[RBD\text{-}CRD{:}PS]$$
$$d[RBD\text{-}CRD{:}PS]/dt = k_3[RAS{:}RBD\text{-}CRD{:}PS] - k_{-3}[RAS][RBD\text{-}CRD{:}PS] - k_4[RBD\text{-}CRD{:}PS]$$
$$d[RAS{:}RBD\text{-}CRD{:}PS]/dt = -k_3[RAS{:}RBD\text{-}CRD{:}PS] + k_{-3}[RAS][RBD\text{-}CRD{:}PS]$$

Three kinetic constants—$k_3$, $k_{-3}$ and $k_4$—are involved. We first estimated $k_3$ by fitting the competitive FRET dissociation assay data. These assays monitor only the dissociation of RAS from the RAS:RBD-CRD:PS complex while preventing the reverse reaction by including an excess of unlabeled RBD-CRD, which sequesters free RAS. Under these conditions, the dissociation of RAS can be treated as an irreversible unimolecular reaction:

RAS:RBD-CRD:PS(m) → RAS(m) + RBD-CRD:PS(m) ($k_3$ fwd rate constant)

By fitting the FRET dissociation curves obtained from TIRF measurements, we estimated $k_3$ to be 0.18 $s^{-1}$ (**Supplementary Fig. 10**). This value corresponds to the unimolecular rate constant for RAS dissociation from the membrane-bound RAS:RBD-CRD:PS complex.

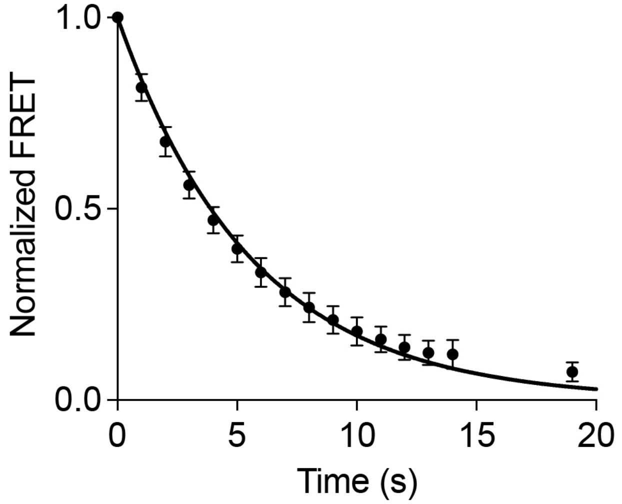


**Supplementary Fig. 10 Competitive FRET dissociation curve fit.** The data was fit with an irreversible unimolecular dissociation kinetic model: $y(t) = e^{-k_3 t}$. $k_3$ was estimated to be 0.18 $s^{-1}$ The experiments were conducted on membranes composed of 77% DOPC, 20% DOPS and 3% MCC-DOPE. Ras density is ~500 $\mu m^{-2}$.

The rate constants $k_{-3}$ and $k_4$ were determined by fitting RBD-CRD free dissociation data obtained at different RAS surface densities. Under these conditions, the membrane-bound forms of RBD-CRD are RAS:RBD-CRD:PS and RBD-CRD:PS. The sum of these two species were optimized to estimate $k_{-3}$ and $k_4$. Except for RBD-CRD that dissociates into solution, all species involved in the reaction remain membrane-bound. Because free RBD-CRD escapes the system due to continuous buffer rinsing, the amounts of all membrane-bound species can be expressed in terms of their surface density ($\mu m^{-2}$). For each dissociation assay, we experimentally determined the surface densities of RAS and membrane-bound RBD-CRD. The assays were performed at 20 nM RBD-CRD which barely saturate RAS. Therefore, we assume that RBD-CRD exists predominantly as RAS:RBD-CRD:PS unless its density exceeds that of RAS. This assumption is supported by data showing that more than 100 nM RBD-CRD is required to observe a substantial amount of RAS-free RBD-CRD (Fig. 1f). Based on our experimental measurements, the following initial conditions were used for curve fitting:

| **Total Ras ($\mu m^{-2}$)** | **RAS:RBD-CRD:PS ($\mu m^{-2}$)** | **RBD-CRD:PS ($\mu m^{-2}$)** |
|---|---|---|
| 1132 | 1132 | 47 |
| 710 | 710 | 8 |
| 383 | 383 | 39 |
| 207 | 207 | 4 |
| 141 | 135 | 0 |

**Supplementary Table 1. Surface densities of membrane-bound species used for the initial conditions of RBD-CRD free dissociation curve fitting.**

The following table summarizes the estimated kinetic constants:

| **Reaction** | **Forward rate constant** | **Reverse rate constant** |
|---|---|---|
| RAS:RBD-CRD:PS(m) ↔ RAS(m) + RBD-CRD:PS(m) | $k_3 = 0.18\ s^{-1}$ | $k_{-3} = 0.043\ \mu m^2 s^{-1}$ |
| RBD-CRD:PS(m) → RBD-CRD(s) + PS(m) | $k_4 = 0.42\ s^{-1}$ | N/A |

**Supplementary Table 2. Estimated kinetic constants for RBD-CRD dissociation**

We used the kinetic constants shown in Supplementary Table 2 for kinetic simulations in Figure 5c. Kinetic modeling and kinetic constant estimation were performed in SimBiology (Mathworks) by numerically solving the rate equations using ODE45.
